# *De-novo* chromosome level assembly of plant genomes from long read sequence data

**DOI:** 10.1101/2021.09.09.459704

**Authors:** Priyanka Sharma, Ardashir Kharabian Masouleh, Bruce Topp, Agnelo Furtado, Robert J. Henry

## Abstract

Recent advances in the sequencing and assembly of plant genomes have allowed the generation of genomes with increasing contiguity and sequence accuracy. The chromosome level assembly of the contigs generated from long read sequencing has involved the use of proximity analysis (Hi-C) or traditional genetic maps to guide the placement of sequence contigs within chromosomes. The development of highly accurate long reads by repeated sequencing of circularized DNA (PacBio HiFi) has greatly increased the size of contigs. We now report the use of HiFiasm to assemble the genome of *Macadamia jansenii*. a genome that has been used as model to test sequencing and assembly. This achieved almost complete chromosome level assembly from the sequence data alone without the need for higher level chromosome map information. Eight of the 14 chromosomes were represented by a single large contig and the other 6 assembled into 2-4 main contigs. The small number of chromosome breaks appear to be due to highly repetitive regions of ribosomal genes that cannot be assembled by these approaches. *De novo* assembly of near complete chromosome level plant genomes now seems possible using these sequencing and assembly tools. Further targeted strategies might allow these remaining gaps to be closed.

**Significance statement (of up to two sentences):** *De novo* assembly of near complete chromosome level plant genomes is now possible using current long read sequencing and assembly tools.

## Introduction

Reference genome sequences are a key resource for plant science. The challenge of producing a complete genome sequence has been greatly reduced by advances in both DNA sequencing (Levy and Myers, 2016; Hon et al., 2020) and sequence assembly tools (Phillippy, 2017; Chen et al., 2017). Final assembly of chromosome level genomes has relied upon evidence other than the sequence data alone such as genetic maps (Fierst, 2015; Yu et al., 2019).

Advancements in the field of sequencing, assembly and scaffolding technologies, along with the rapid increase in the freely available genomic data (https://www.ncbi.nlm.nih.gov/genbank/statistics/), has greatly facilitated the development of highly accurate *de novo* assemblers.

Short-read *de novo* assemblers are not efficient in assembling the complex and long repetitive regions of plant genomes, like centromere and telomere (Liao et al., 2019). To address this limitation, long read sequencing technologies, also known as third generation sequencers, have been developed. However, these long reads from PacBio and Oxford Nanopore have been less accurate with an average of 90% accuracy as compared to the 99.9% accuracy of Illumina reads (Amarasinghe et al., 2020; Shendure et al., 2017). Therefore, hybrid assembly pipelines were used to assemble a large number of genomes, to overcome the shortcomings of both the long reads and short reads. This has allowed assembly of larger contigs from complex genomes. However, to achieve chromosome level genome assembly, scaffolding of the contigs is usually required. Analysis of sequence proximity in the chromatin by methods such as Hi-C have made this possible(Dudchenko et al., 2017).

Recent advancements in long read sequencing technology have allowed a single molecule to be sequenced multiple times to produce long high fidelity reads (Pac Bio HiFi) with base level accuracy of 99.9 % (Wenger et al., 2019b).

*Macadamia jansenii* has been used to compare methods for the sequencing and assembly of plant genomes (Murigneux et al., 2020; Sharma et al., 2021a). This genome has size (∼ 800Mb) typical of many plant genomes but with a relatively low heterozygosity (Sharma et al., 2020b). Assembly of this genome using highly accurate circular consensus reads (CCS) reads (Pac Bio HiFi) was found to give a more contiguous genome than that obtained with earlier longer continuous long reads (Pac Bio CLR) (Sharma et al., 2021a). HiFiasm has been used to successfully assemble Fragaria *× ananassa* (garden strawberry), *Rana muscosa* (mountain yellow-legged frog) and *Sequoia sempervirens* (California redwood) (Cheng et al., 2021). Recently HiFiasm was reported to allow highly contiguous assembly of plant genomes (Druiguez et al., 2021). We now report the near complete chromosome level assembly of the *M. jansenii* genome from HiFi reads with the HiFiasm assembly tool and an analysis of the assembled genome against a HI-C chromosome level assembly.

## Results

### HiFiasm assembly

The estimated genome size of *M. jansenii* is 780 Mb (Murigneux et al., 2020). The size of the primary Hifiasm assembly was 826 Mb including 779 contigs **(Table S1)**, with the longest contig of 71.9 Mb and an average contig length of 1 Mb. BUSCO analysis showed that the assembly covered 99.6% of universal single copy genes (**Table 1**). The contigs generated in this assembly were characterized in three groups based upon their size, large contigs > 1 Mb; medium size contigs, between 1 Mb and. 100 Kb and small contigs < 100 Kb.

**Table 1:** (A) HiFiasm contigs in different size categories. (B) Comparison of Primary and Haploid assemblies generated from HiFiasm genome assembler tool.

| (A) HiFiasm Assembly | Number of contigs | Assembly length | N50 | N75 | BUSCO |
| --- | --- | --- | --- | --- | --- |
| Total Contigs | 779 | 826 Mb | 46 Mb | 25 Mb | 99.60% |
| Contigs >10 Mb | 19 | 746 Mb | 48 Mb | 39 Mb | 93.90% |
| Contigs > 42 Mb | 23 | 772 Mb | 48 Mb | 30 Mb | 99.10% |
| Contigs >1 Mb | 30 | 784 Mb | 46 Mb | 30 Mb | 99.10% |
| Contigs > 100 Kb | 94 | 805 Mb | 46 Mb | 27 Mb | 99.00% |
| Between <1 Mb to 100kb | 64 | 20 Mb | 0.49 Mb | 0.22 Mb | 0.20% |
| Between <100 Kb to 10 Kb | 685 | 22 Mb | 0.032 Mb | 0.028 Mb | 0.00% |

| (B) HiFiasm Primary and Haploid assemblies' comparison |  |  |  |  |  |
| --- | --- | --- | --- | --- | --- |
| Primary assembly | 779 | 827 Mb | 46.1 Mb | 25 Mb | 99.60% |
| Hap 1_assembly | 879 | 816 Mb | 24.4 Mb | 8.9 Mb | 98.80% |
| Hap 2_assembly | 363 | 776 Mb | 14.3 Mb | 5.4 Mb | 97.90% |
| Hap 1 > 1 Mb | 96 | 736 Mb | 16.4 Mb | 6.8 Mb | 96.70% |
| Hap 2 > 1Mb | 72 | 766 Mb | 24.5 Mb | 12.3 Mb | 98.10% |

#### Larger size contigs > 1 Mb

There were 30 contigs greater than 1Mb in length. These contigs alone provided a good assembly with an N50 of 46 Mb and a BUSCO score of 99.1% **(Table 1)**. Dotplot analysis against the Hi-C assembly showed that of the 9 contigs more than 46 Mb in length, 8 correspond to complete Hi-C pseudomolecules (i.e., each contig corresponds to a single chromosome (chromosome # 1, 4, 5, 6, 10, 11, 13 &14) **(Figure 2 (A)**. One contig (Ptg000010|), corresponded to a large part of the second largest chromosome (chromosome 2). Two other contigs of about 25 Mb and 2.7 Mb covered the other parts of this chromosome **(Table 2 & 3), (Figure 2 (B))**. The 14 contigs between 4 Mb and 46 Mb in size, covered the remaining six chromosomes, in combinations of two to four contigs. Five of the contigs between 1 Mb and 4Mb in size, corresponded to nuclear ribosomal RNA sequences, and the other two contigs matched parts of chromosome 2 and 7 **(Table 2 & 3, Figure 2 (B))**.

**Table 2:** Chromosomal location of HiFiasm contigs > 1 Mb.

| Contig id > 1 Mb | Length in bp | Hi-C pseudo-molecule corresponding<br>HiFiasm contigs |
| --- | --- | --- |
| ptg000016l | 71935981 | Chr 1 + Ribo RNA |
| ptg000003l | 57251071 | Chr 6 |
| ptg000017l | 57081251 | Chr 4 |
| ptg000011l | 56513637 | Chr 5 |
| ptg000004l | 49863231 | Chr 10 |
| ptg000012l | 48320516 | Chr 11 |
| ptg000023l | 47997562 | Chr 13 |
| ptg000010l | 46138073 | Chr 2 + Ribo RNA |
| ptg000008l | 46131124 | Chr 14 |
| ptg000014l | 43049961 | Chr 9 |
| ptg000009l | 39279660 | Chr 3 |
| ptg000002l | 29700554 | Chr 8 |
| ptg000001l | 26771894 | Chr 12 |
| ptg000006l | 25189511 | Chr 2 |
| ptg000007l | 23138637 | Chr 7 + Ribo RNA |
| ptg000013l | 22539440 | Chr 8 |
| ptg000020l | 22399594 | Chr 7 |
| ptg000052l | 20335125 | Chr 12 |
| ptg000021l | 13354688 | Chr 3 |
| ptg000019l | 8098418 | Chr 7 |
| ptg000022l | 6676624 | Chr 3 |
| ptg000005l | 6127021 | Chr 9 |
| ptg000072l | 4271045 | Chr 12 |
| ptg000018l | 2743534 | Ribo RNA |
| ptg000025l | 2713795 | Part of Chr 2 |
| ptg000062l | 1651603 | Ribo RNA |
| ptg000074l | 1299006 | Ribo RNA |
| ptg000034l | 1171806 | Ribo RNA |
| ptg000036l | 1154310 | Ribo RNA |
| ptg000033l | 1122141 | Part of Chr 7 |

**Table 3:** Hifiasm contigs (> 1 Mb) covering each of the Hi-C pseudo-molecules.

| *M. jansinii*<br>HiC pseudo-molecules<br>(A) | Size of HiC Pseudo-molecules<br>(B) | Hifiasm Contigs Corresponding to HiC<br>scaffolds<br>(C) | Hifiasm contigs length (with explanation)<br>(D) | Hifiasm Combined contigs<br>length<br>(E) | Extra Hifiasm<br>length (Hifiasm<br>contig length - HiC<br>scaffold length)<br>(E-B) |
| --- | --- | --- | --- | --- | --- |
| Chr 1 | 67682215 | ptg000016l | 71935981 | 71935981 | 4253766 |
| Chr 2 | 63669590 | ptg000006l + ptg000025l + ptg000010l | 74041379 (= 25189511 + 2713795 + 46138073) | 74041379 | 10371789 |
| Chr 3 | 58143993 | ptg000021l + ptg000009l + ptg000022l | 59310972 (= 13354688 + 39279660 + 6676624) | 59310972 | 1166979 |
| Chr 4 | 56076407 | ptg000017l | 57081251 | 57081251 | 1004844 |
| Chr 5 | 55220784 | ptg000011l | 56513637 | 56513637 | 1292853 |
| Chr 6 | 53595462 | ptg000003l | 57251071 | 57251071 | 3655609 |
| Chr 7 | 52077970 | ptg000019l + ptg000020l + ptg000033l +<br>ptg000007l | 54758790 (= 8098418 + 22399594 + 1122141 +<br>23138637) | 54758790 | 2680820 |
| Chr 8 | 49563658 | ptg000013l + ptg000002l | 52239994 (= 22539440 + 29700554) | 52239994 | 2676336 |
| Chr 9 | 49085581 | ptg000014l + ptg000005 | 49176982 (= 43049961 + 6127021) | 49176982 | 91401 |
| Chr 10 | 48974653 | ptg000004l | 49863231 | 49863231 | 888578 |
| Chr 11 | 47698009 | ptg000012l | 48320516 | 48320516 | 622507 |
| Chr 12 | 46713600 | ptg000001l + ptg000072l + ptg000052l | 51378064 (= 26771894 + 4271045 + 20335125) | 51378064 | 4664464 |
| Chr 13 | 45610911 | ptg000023l | 47997562 | 47997562 | 2386651 |
| Chr 14 | 45288529 | ptg000008l | 46131124 | 46131124 | 842595 |

**Figure 1:**
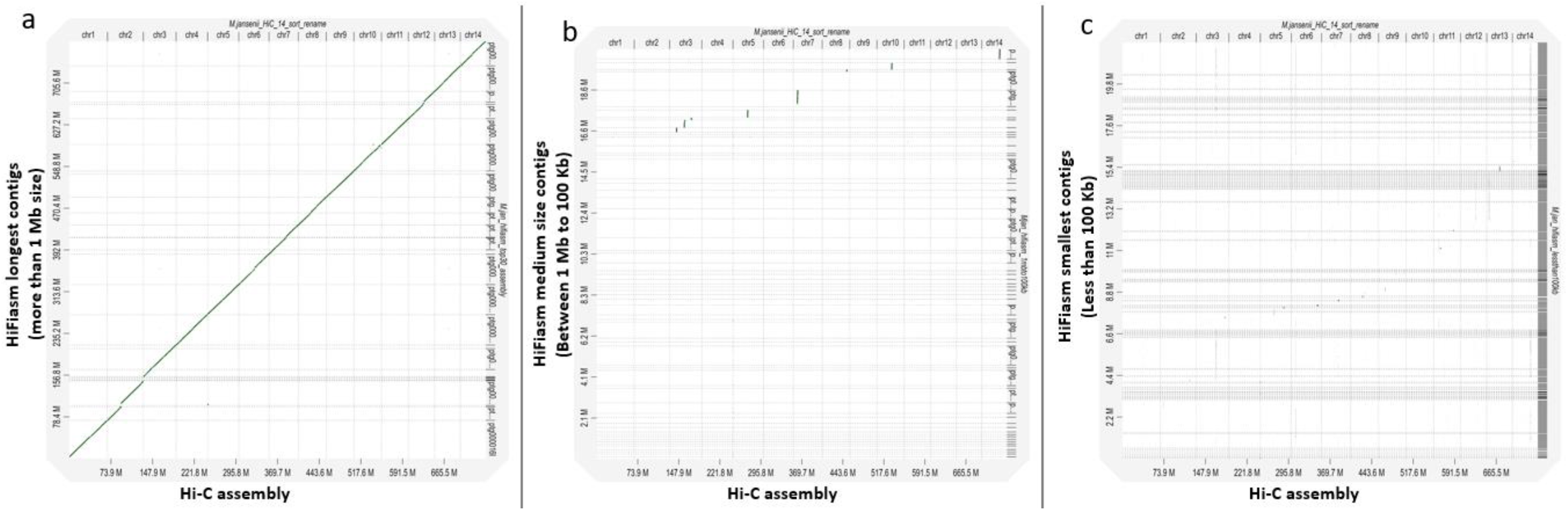
Dotplot of *M. jansenii* Hi-C genome assembly against HiFiasm contigs. (a) HiFiasm longest contigs (> 1 Mb size). (b) HiFiasm medium size contigs (< 1 Mb and > 100 Kb) and (c) HiFiasm smallest contigs (<100 Kb).

**Figure 2:**
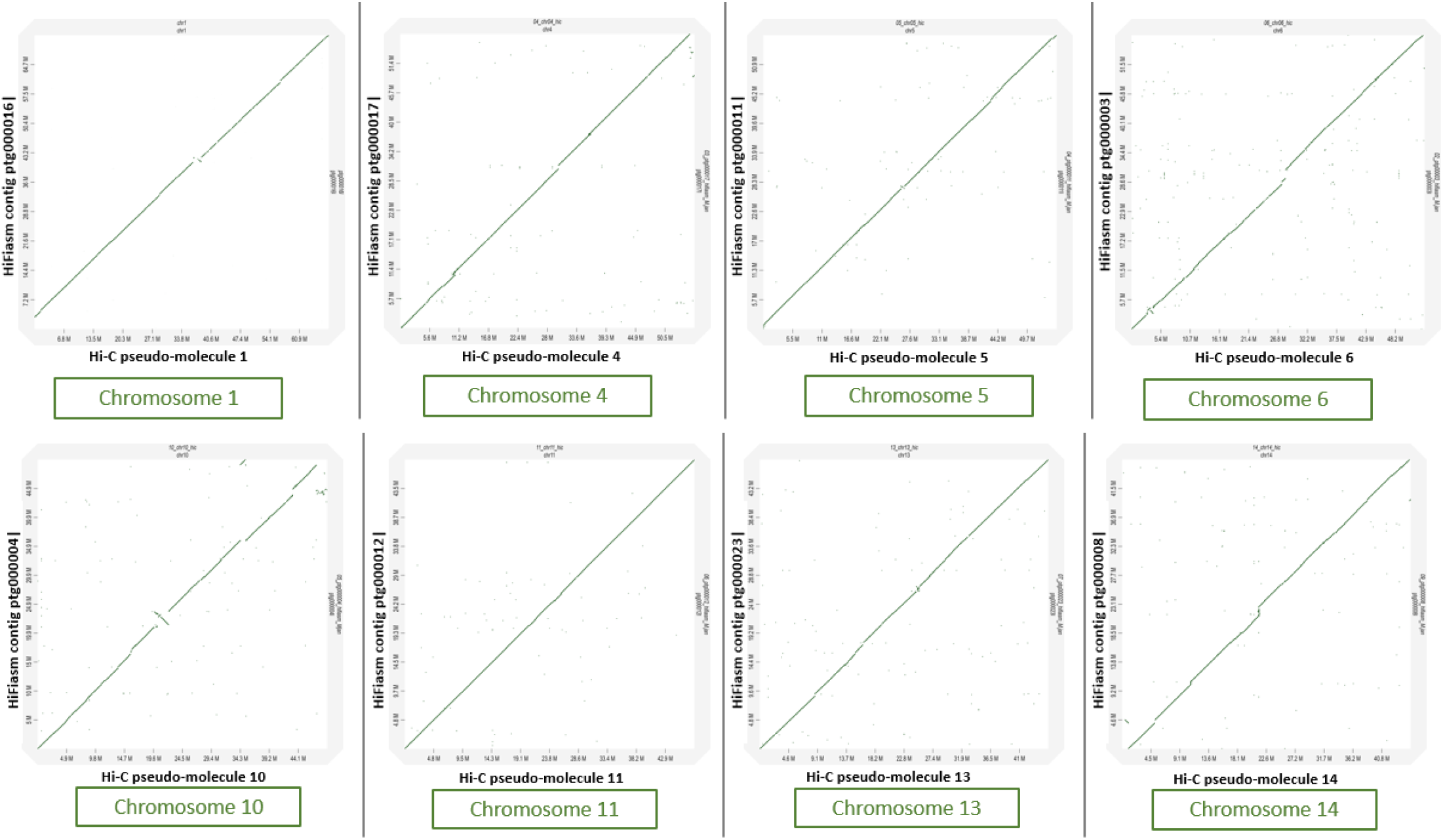
Dotplots of HiFiasm contigs against Hi-C pseudo-molecules. A) Pseudo-molecules which are covered by single HiFiasm contig. B) Pseudo-molecules which are covered by more than one HiFiasm contig.

**Figure 2(B):**
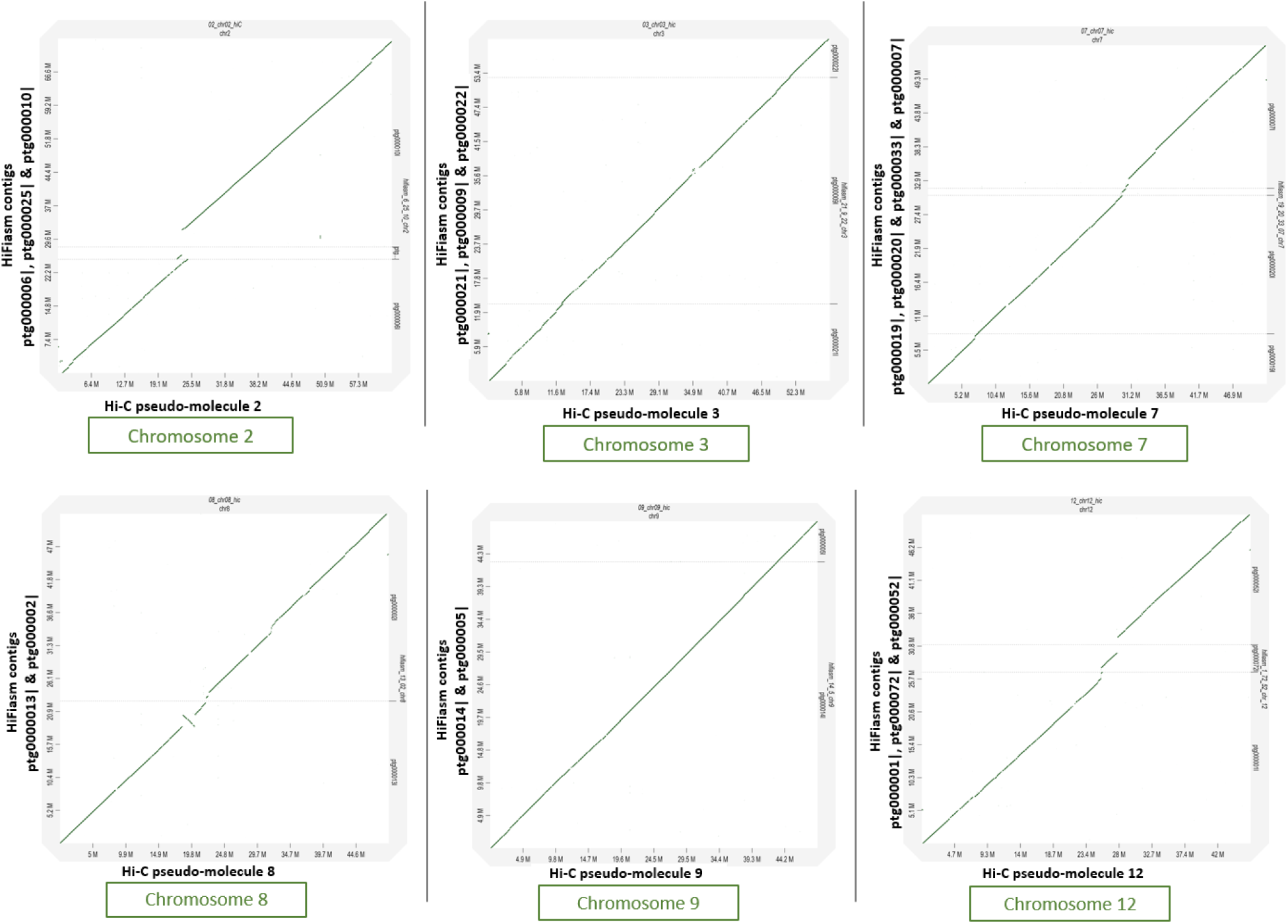
Pseudo-molecules which are covered by more than one HiFiasm contig.

#### Medium size contigs

There were 64 contigs between 1Mb and 100 Kb in size. These contigs had 0% BUSCO genes **(Table 2)**. Only eight contigs in the range between100 Kb and 824 Kb corresponded to seven Hi-C pseudo-molecules (with an alignment block length of more than 100Kb) **(Figure 1B, Table S 2, & Figure S2, S3, Table S2 and S3)**. Out of these eight contigs, five contigs corresponds to the terminal part of the Hi-C pseudo-molecules and three contigs corresponds to the non-terminal regions of Hi-C chromosomes 3 and 7, marked as red starts in **Figure S2 (A) & (B)**. The majority of the medium size contigs corresponds to ribosomal RNA genes **(Figure 5 B)** and one contig of 183 Kb corresponded to a chloroplast assembly **(Figure 3 (B))**. None of the contigs showed similarity with mitochondrial sequences **(Figure 4 B)**.

**Figure 3:**
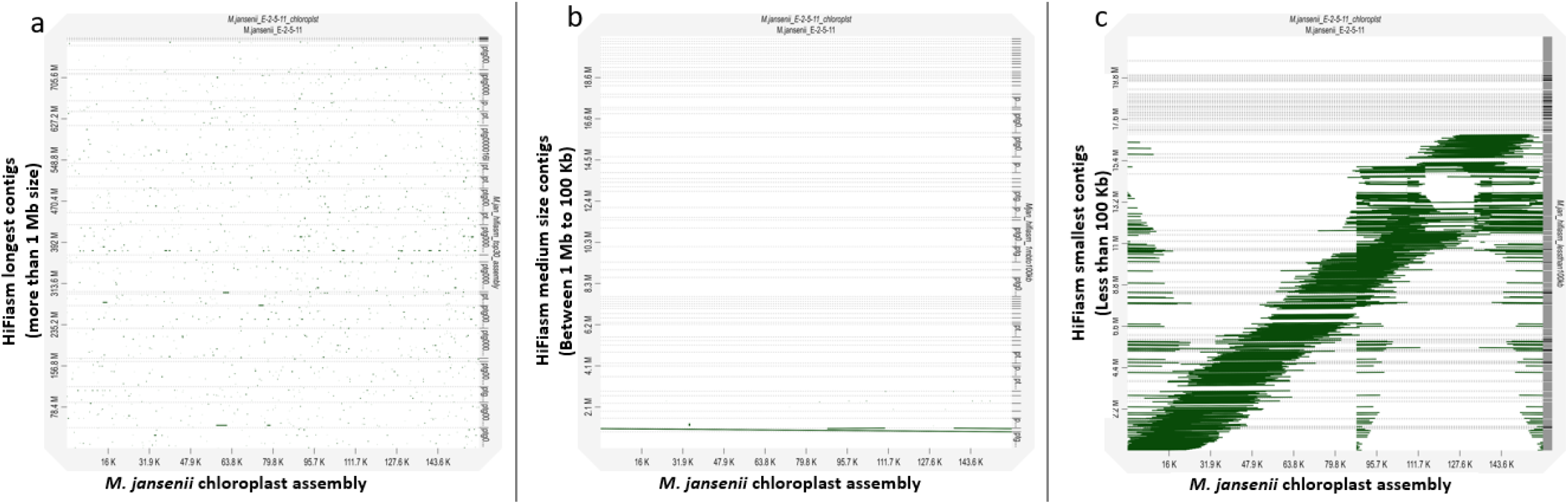
Dotplot of *M. jansenii* chloroplast genome sequence against HiFiasm contigs. (a) HiFiasm longest contigs (> 1 Mb size). (b) HiFiasm medium size contigs (< 1 Mb and > 100 Kb) and (c) HiFiasm smallest contigs (<100 Kb).

**Figure 4:**
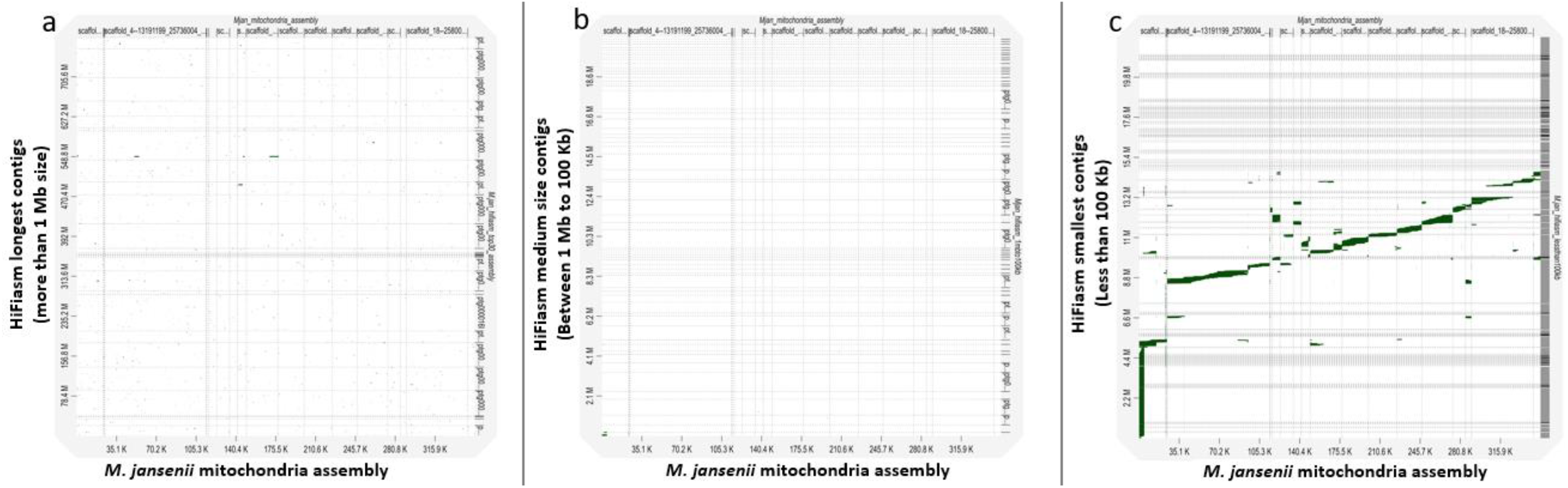
Dotplot of *M. jansenii* mitochondria genome sequence against three sets of HiFiasm contigs. (a) HiFiasm longest contigs (> 1 Mb size). (b) HiFiasm medium size contigs (< 1 Mb and > 100 Kb) and (c) HiFiasm smallest contigs (<100 Kb).

**Figure 5:**
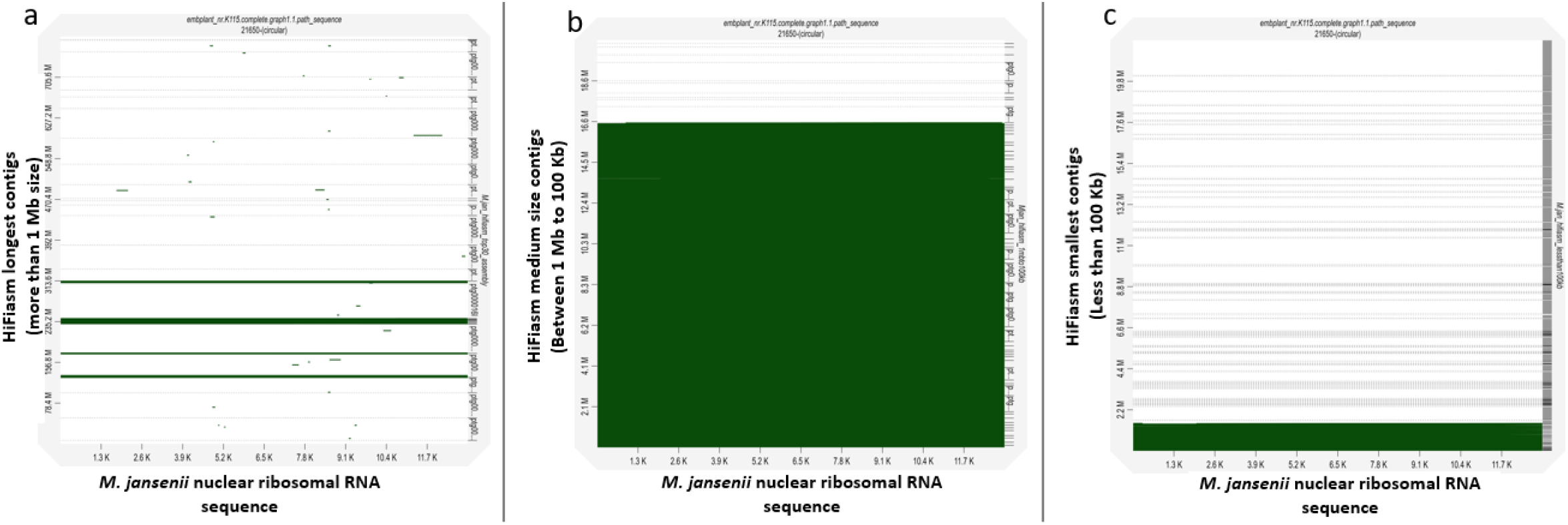
Dotplot of *M. jansenii* nuclear ribosomal RNA sequence against HiFiasm contigs. (a) HiFiasm longest contigs (> 1 Mb size). (b) HiFiasm medium size contigs (< 1 Mb and > 100 Kb) and (c) HiFiasm smallest contigs (<100 Kb).

#### Smaller contigs

There were 685 contigs between 10 Kb and 100 Kb in size. The majority of these small size contigs from the HiFiasm assembly corresponded to small portions of the chloroplast and mitochondrial genomes. These contigs aligned together covered the complete organelle genomes **(Figure 3 (C), 4 (C))**. However, a few of contigs corresponded to nuclear ribosomal RNA sequences **(Figure 5 (C)**. This contig set also showed 0% BUSCO genes **(Table 1)**.

### Influence of data volume

HiFiasm assembly from CCS reads from two individual SMRT cells and the combined data is given in (**Table S4)**. A HiFiasm assembly generated from the 10 X CCS data produce 4511 contigs with 909 Mb of assembly length and N50 of 0.38 Mb, whereas a larger CCS file with 18 X coverage generated an assembly with less contigs (1058), a shorter assembly length (833 Mb) with an improved N50 of 4.4 Mb **(Table 1)**. The 18 X assembly closer to the combined CCS assembly (and Hi-C assembly), than the 10X assembly.

Haploid assembly details are given in **Table 1**. The haploid 1 assembly has a greater number of contigs than the haploid 2 assembly. BUSCO results were similar for the two haploid and primary assemblies as all assemblies were relatively complete.

### Comparison with Hi-C assembly

A dotplot analysis of 14 pseudo-molecules of *M. jansenii* Hi-C assembly against the HiFiasm assembly is shown in **Figure 1**. The dotplot of contigs >1Mb in size showed a complete match of 25 contigs (out of total 30) with the 14 Hi-C pseudo-molecules (Sharma et al., 2021b) (**Figure 1(A)**). The remaining five large contigs did not contribute to the genome assembly. They were composed of nuclear ribosomal RNA sequences. Chromosomes 1, 4, 5, 6, 10, 11, 13 and 14 were covered by single contig of the HiFiasm assembly **(Figure 2 (A)**, two chromosomes (Chr 8 and 9) were covered by two contigs and chromosomes 2, 3 and 12 were covered by three contigs and chromosome seven by four contigs **(Figure 2(B) & Table 2 & 3)**.

### Organelle genome analysis

Dotplot analysis of a 159 Mb full length chloroplast genome against the HiFiasm genome assembly indicated the insertion of small fragments of chloroplast sequences in the nuclear genome assembly **(Figure 3 (A), Figure S1 (A))**, which also align with the Hi-C assembly results (Sharma et al., 2021b) **(Figure S1 (B)**. Secondly, among the middle size contig set, only one contig (ptg0000186|) of 183 Mb aligned with the chloroplast genome **(Figure 3 (B)**. Contig ptg000186| covered the complete chloroplast genome including the two inverted repeat regions of the chloroplast (**Figure S4)**. Another HiFiasm middle size contig, ptg000066|, also showed some similarity with the chloroplast assembly and also aligned with the terminal end of Hi-C chromosome 14 **(Figure S5)**. Analysis of the smaller size contigs showed the majority of these contigs contains some fragments of the chloroplast assembly **(Figure 3 (C))**.

Mitochondrial sequence analysis was performed. The size of the *de-novo* mitochondrial assembly was 351 Kb. Analysis against the HiFiasm assembly, indicated the presence of mitochondrial sequences in the smallest set of contigs. The majority of these contigs cover small fragments of the mitochondria genome **(Figure 4 (C)**. Whereas, in the larger contig set, (>1 Mb), only a few contigs showed some similarity with mitochondrial sequences. These represent the mitochondria sequences inserted in the nuclear the genome **(Figure 4 (A)**, which aligns with the dotplot result of Hi-C assembly **(Figure S1 (B) b)**. The middle size contigs did not show the presence of any mitochondria sequences in the dotplot analysis **(Figure 4 (B))**.

### Nuclear ribosomal RNA gene sequences analysis

Dotplot analysis of nuclear ribosomal RNAs sequences showed matches with the majority of the middle size contigs with a small number of contigs from the smaller set of contigs having ribosomal RNAs sequence **(Figure 5 (b) & (c))**.

### Analysis of repeat elements

The Hifiasm assemblies were larger than the corresponding Hi-C pseudomolecules **(Table 3)**. This is probably because the HiFiasm contigs included a larger proportion of repetitive elements than the corresponding Hi-C pseudomolecules **(Table 4)**. The longer chromosome had a generally higher content of repetitive elements suggesting that the presence of these repeat regions explained their greater size.

**Table 4:** Comparative repetitive elements of Hi-C pseudo-molecules and HiFiasm assembly

| *M. jansinii*<br>pseudo-molecules | Genome assembler | Size of Pseudo-molecules | Total repeats | LINE | LTR | DNA elements | Unclassified | Simple repeats |
| --- | --- | --- | --- | --- | --- | --- | --- | --- |
| Chr 1 | Hi-C | 67682215 | 62% | 4.13% | 30.25% | 0.52% | 26.80% | 0.64% |
|  | HiFiasm | 71935981 | 62.19% | 3.88% | 34.78% | 0.58% | 22.64% | 0.65% |
| Chr 2 | Hi-C | 63669590 | 66% | 3.31% | 38.21% | 1.12% | 23.34% | 0.86% |
|  | HiFiasm | 74041379 | 68% | 2.98% | 39.58% | 1.22% | 23.78% | 0.44% |
| Chr 3 | Hi-C | 58143993 | 52.34% | 6.15% | 20.48% | 1.13% | 24.18% | 0.67% |
|  | HiFiasm | 59310972 | 54.30% | 7.93% | 20.70% | 0.97% | 24.09% | 0.72% |
| Chr 4 | Hi-C | 56076407 | 55.07% | 6.26% | 22.79% | 0.79% | 24.27% | 1.13% |
|  | HiFiasm | 57081251 | 57.20% | 7.21 % | 21.57 % | 0.90 % | 26.58 % | 1.23 % |
| Chr 5 | Hi-C | 55220784 | 53.10% | 3.27% | 31.43% | 0.96% | 16.99% | 0.78% |
|  | HiFiasm | 56513637 | 54.29 % | 3.47 % | 31.55 % | 0.75 % | 17.93 % | 0.85 % |
| Chr 6 | Hi-C | 53595462 | 55.10% | 8.75% | 19.94% | 1.18% | 25.49% | 0.79% |
|  | HiFiasm | 57251071 | 58.78 % | 9.47 % | 22.60 % | 1.43 % | 24.01 % | 1.44 % |
| Chr 7 | Hi-C | 52077970 | 52.89% | 7.04% | 21.33% | 1.42% | 22.38% | 0.85% |
|  | HiFiasm | 54758790 | 51.21 % | 6.65 % | 21.25 % | 1.25 % | 21.37 % | 0.79 % |
| Chr 8 | Hi-C | 49563658 | 43.97% | 5.39% | 15.47% | 0.64% | 21.29% | 1.25% |
|  | HiFiasm | 52239994 | 41.81% | 6.25% | 21.97% | 1.30% | 12.29% | 0% |
| Chr 9 | Hi-C | 49085581 | 48.13% | 5.39% | 18.38% | 1.76% | 21.97% | 0.86% |
|  | HiFiasm | 49176982 | 45.93 | 5.41% | 24.84% | 1.65% | 14.03% | 0% |
| Chr 10 | Hi-C | 48974653 | 48.10% | 6.24% | 17.30% | 0.90% | 22.83% | 1.02% |
|  | HiFiasm | 49863231 | 44.66% | 5.91% | 22.85% | 2.82% | 13.08% | 0% |
| Chr 11 | Hi-C | 47698009 | 48.10% | 6.24% | 17.30% | 0.90% | 22.83% | 1.02% |
|  | HiFiasm | 48320516 | 44.70% | 4.01% | 26.25% | 2.82% | 11.62% | 0% |
| Chr 12 | Hi-C | 46713600 | 44.35% | 4.47% | 16.46% | 1.39% | 21.29% | 0.85% |
|  | HiFiasm | 51378064 | 25.64 | 4.47% | 21.17% | 2.44% | 11.73% | 0% |
| Chr 13 | Hi-C | 45610911 | 42.22% | 5.52% | 13.67% | 0.70% | 21.14% | 1.31% |
|  | HiFiasm | 47997562 | 39.68% | 5.42% | 18.80% | 1.63% | 13.76% | 0% |
| Chr 14 | Hi-C | 45288529 | 42.53% | 5.82% | 12.85% | 0.96% | 21.53% | 1.50% |
|  | HiFiasm | 46131124 | 41.10% | 5.53% | 19.60% | 1.94% | 13.92% | 0% |

## Discussion

This era of genomics is continuing to advance with improved sequencing technologies and the potential to sequence all the species on earth (Lewin et al., 2018).

Accurate chromosome level genome assembly requires accurate reads, high genome coverage and long read length. This has typically involved the use of very high coverage and data from multiple sequencing platforms along with mapping of Hi-C technologies to achieve chromosome level assemblies. However, the combination of high sequence accuracy in a long read in HiFi reads (99.8% accuracy at around 15 Kb average length) provides the option to assemble a complete genome using a single sequencing technology (Cheng et al., 2021) and with a more readily obtainable genome coverage (Wenger et al., 2019a).

Here, we have combined the benefit of this highly accurate reads along with an improved assembly tool HiFiasm (Cheng et al., 2021).HiFi read genome coverage of 28-40 X, for plant genomes within the range of 700 −1000 Mb size, was sufficient to generate high quality assemblies with Mb contig sizes (Sharma et al., 2021b). The extracted DNA from *Macadamia jansenii* contained some impurities, as might be expected for many samples. Two SMRT cells were required to generate 28 X genome coverage with CCS reads. For some samples this may be possible with one single run providing the required coverage if sufficient DNA purity is achieved reducing the cost of obtaining sufficient sequence. When the two individual CCS runs of 10 X and 18 X, were assembled separately using HiFiasm, the final assembly was very fragmented (N50 of 0.38 Mb and 4.4 Mb, respectively) for *M. jansenii* (Table S1), whereas the combined 28X gave a highly continuous assembly with N50 of 46.1 Mb and 99.6 % BUSCO results. Suggesting if the DNA isolation methods is highly efficient in removing impurities, then a single run with less coverage would be sufficient to assemble the genome. Which implies, lesser the data volume, so do the computational requirements. The higher base-calling accuracy by HiFi improves the assembly accuracy by bypassing many time consuming and heavy computational requirement steps in the assembly workflow. The *M. jansenii* assembly from HiFiasm using HiFi sequencing data produced a near chromosome level assembly, with eight contigs covering eight complete Hi-C pseudo-molecules and other six chromosomes, being covered by only one to four breaks by total of 17 contigs. The gaps in those six chromosomes were found to be due to the presence of large numbers of ribosomal RNA gene sequences (need to look into Dotplot paf file, with Hi-C and HiFiasm). For a plant with ∼800 Mb data, we estimate a high-quality chromosome level assembly could be produced within a week from the plant material, if the DNA extraction step is well established.

This highly continuous *M. jansenii* chromosome level assembly will help in better understanding of the genome of macadamias. All four species of Macadamia are listed as threatened under Australian legislation (Mast et al., 2008), but *M. jansenii* is particularly endangered, due to its very low population size (less than 200 plants in the wild) (Shapcott and Powell, 2011). The highly accurate genome assembly will facilitate its conservation and use in breeding. *M. jansenii* has small inedible nuts and (Gross and Weston, 1992). However, due to its small tree size and narrow root spread, it is being tested as a rootstock and in hybrids with the commercial species, *M. intergrifolia* (Alam et al., 2018). The HiFiasm assembly (BUSCO 99 %) is much better than the Hi-C assembly (BUSCO 97%) (Sharma et al., 2021b) suggesting the incorporation of some regions missing in the Hi-C assembly.

The initiative to complete the genome assembly of almost all living organisms (Lewin et al., 2018; Koepfli et al., 2015), requires a highly efficient assembly method with sustainable financial, computational and time requirements without compromising on genome accuracy. Continuity and completeness should be taken into consideration (Rhie et al., 2021). Our analysis suggests that HiFiasm assembly with the HiFi reads may require almost no further scaffolding for the plants with similar genome size ∼800 Mb. Analysis of the nature of the few remaining regions of the genome that are not assembled in these analyses may allow the development of targeted strategies to complete these assemblies. Analysis of the sequences at the ends of the contigs formed by HiFiasm assembly of HiFi reads may identify those contigs that have been interrupted by repetitive sequences that cannot be assembled de novo. This technology is successfully assembling regions with high levels of the repeat sequences that make up more than 50% of the *M. jansenii* genome (Sharma et al., 2021b). It may be that the very high accuracy of the HiFi reads detects minor variations in repeat sequences that allow their unique assembly and that only perfect repeats that are longer than the HiFi reads create a barrier to assembly. This study suggests that the large ribosomal gene clusters in the genome of plants may be one of the few limitations to complete assembly. This would suggest that manual analysis could be used to infer that contigs terminating in ribosomal gene repeats might be linked to other contigs ending in ribosomal contigs in the genome. However, additional information may be required for the plants with very large and complex genomes. This approach will be useful, for producing plant genomes, generating high quality *de novo* chromosome level assemblies, especially for laboratories with limited financial, technical and computational resources.

### Experimental procedures

**Sequencing data** Short read (Illumina) sequencing data analysed was from (Murigneux et al., 2020) and long read data (PacBio HiFi) was from (Sharma et al., 2021a).

### Hifiasm Assembly

The HiFiasm genome assembly (Cheng et al., 2021) was generated using High Performance Computing (HPC) facility at University of Queensland. For assembly, 24 core processing units (CPUs) and 120 Gb of memory was employed. Default settings of the HiFiasm assembler were used to assemble heterozygous genomes with built-in duplication purging parameters. HiFiasm output directory consists of two haploid (1 & 2), one primary contig and one alternate haplotig GFA graph files. Each halploid and one primary contig GFA file was converted to FASTA format using awk command.

### Analysis of Assembly

The primary HiFiasm assembly of *M. jansenii* included 779 contigs that were categorised into three subsets a) contigs less than 1Mb size, b) Contigs less than 1Mb and more than 100Kb size and c) contigs less than 100 Kb size. Along with the main primary and two haploid assemblies and all three sets of primary contig subsets were passed through analysis using QUAST (Gurevich et al., 2013), BUSCO (Simão et al., 2015) and RepeatModeler (Humann et al., 2019).

### Comparison with Hi-C assembly

The Hifiasm contigs were compared with the *M. jansenii* 14 pseudo-molecules from the Hi-C assembly (Sharma et al., 2021b) using online interactive dotblot tool (Cabanettes and Klopp, 2018).

### Characterisation of organelle genomes content of Hifiasm contigs

A reference mitochondrial genome, chloroplast genome and nuclear ribosomal RNA sequence from this sample were assembled from Illumina raw reads (Murigneux et al., 2020) using the GetOrganelle (Jin et al., 2020) with default parameters. The HiFiasm contigs (779) were compared with the organellar and ribosomal sequences in dotblots.

### Accession numbers

The Hifiasm assembly, chloroplast assembly, mitochondria assembly and nuclear ribosomal RNA sequence of *M. jansenii* has been deposited under NCBI bioproject PRJNA694456.

## Acknowledgements

This project was funded by the Hort Frontiers Advanced Production Systems Fund as part of the Hort Frontiers strategic partnership initiative developed by Hort Innovation, with co-investment from The University of Queensland, and contributions from the Australian Government. We thank the Research Computing Centre (RCC), University of Queensland for support and providing high performance computing resources.

## Short legends for Supporting Information

Figure S1: Comparison of HiFiasm and Hi-C pseudo-molecules against the Chloroplast, mitochondria and nuclear ribosomal RNAs.

Figure S2: (a) Dotplots of Hi-C pseudo-molecules against HiFiasm contigs (Longest contigs >1Mb). (b) Dotplots of Hi-C pseudo-molecules against HiFiasm contigs (Longest and Middle size contigs).

Figure S3: (a) Dotplots of Hi-C pseudo-molecules against HiFiasm contigs (Longest contigs >1Mb). S2 (b) Dotplots of Hi-C pseudo-molecules against HiFiasm contigs (Longest and Middle size contigs).

Figure S4: Chloroplast assembly covered by a single HiFiasm Contig (Ptg0000186|) and small parts by Ptg000066|.

Figure S5: Chloroplast sequence (Ptg0000186| & Ptg000066|) insertions in the Hi-C assembly

Table S1: HiFiasm contigs

Table S2: Middle size HiFiasm contigs (< 1Mb and > 100 Kb) which are part of Hi-C pseudo-molecule assembly.

Table S3: HiFiasm contigs (biggest contigs & middle size contigs) corresponding to *M. jansenii* Hi-C 14 pseudo-molecules.

Table S4: IPA and HiFiasm assembly details

## Author contributions

Contributions of authors were as follows: Designed the study and supervised the project: RJH, AF, AM and BT. Genome assembly and analysis: PS, AM. Data analysis: PS, AF, AM & RJH. Tables and Figures: PS, RJH, and AF. Drafted the manuscript: PS and RJH. Data deposition: PS. All authors edited and approved the final manuscript.

