## Supplementary file for "*De-novo* chromosome level assembly of plant genomes from long read sequence data"

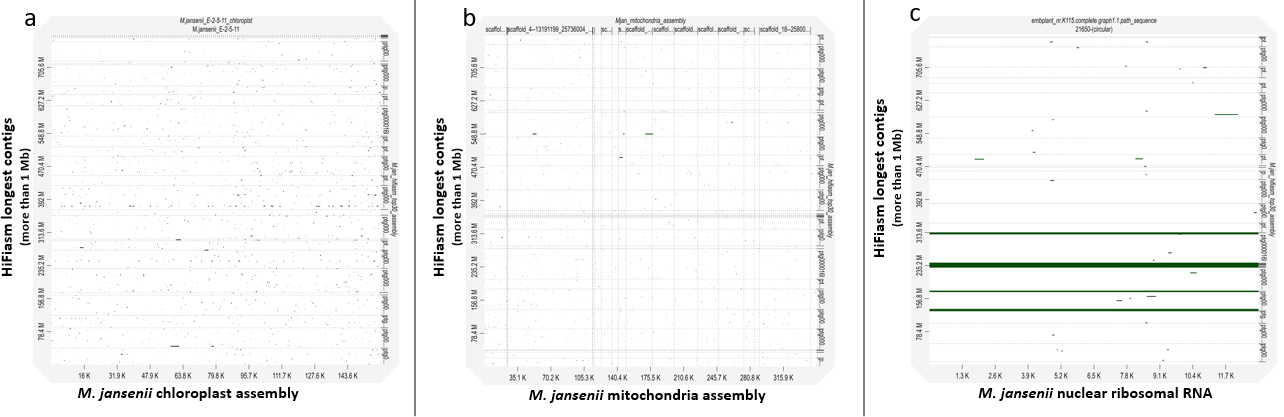

**Figure S1 (A):** Dotplot of *M. jansenii* Hifiasm longest contigs (more than 1 Mb) against a) Chloroplast, b) Mitochondria and c) Nuclear ribosomal RNA sequence of *M. jansenii*.

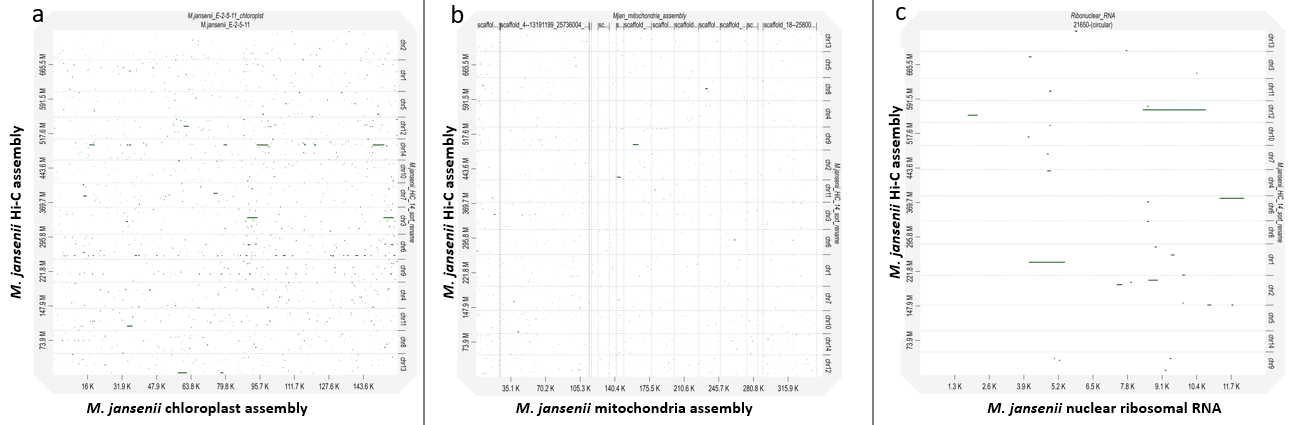

**Figure S1 (B)** Dotplot of *M. jansenii* Hi-C assembly against a) Chloroplast, b) Mitochondria and c) Nuclear ribosomal RNA sequence of *M. jansenii*.

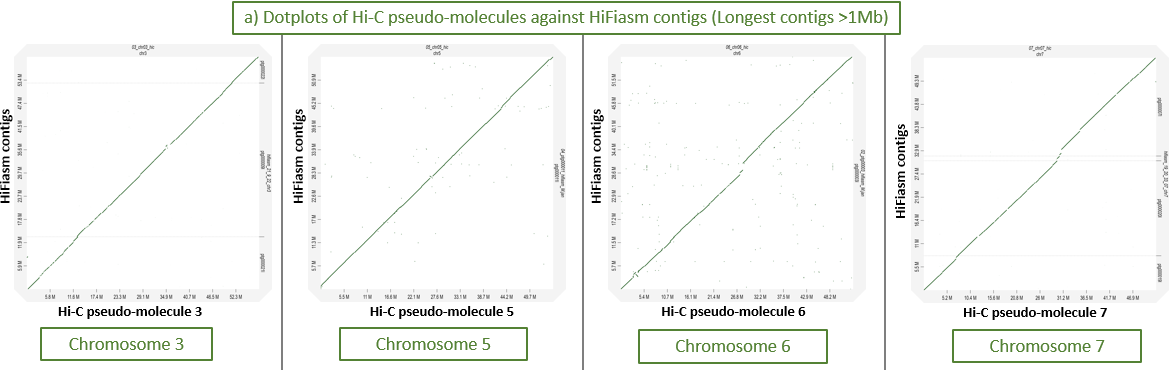

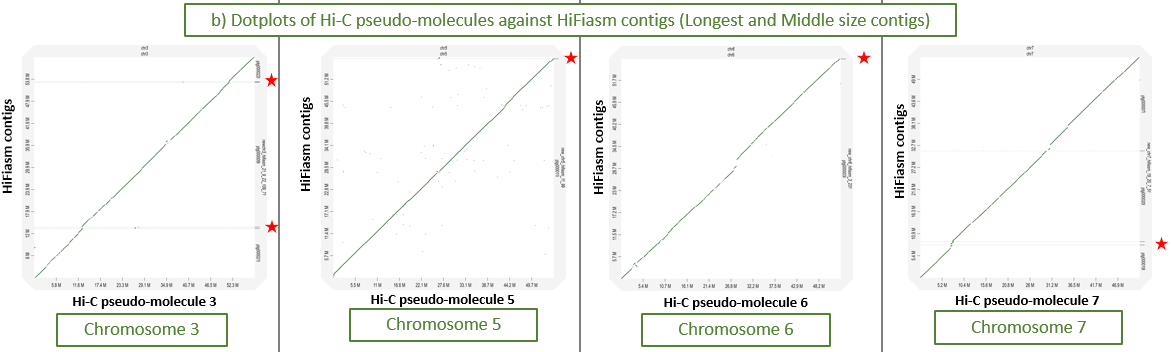

**Figure S2:** (a) Dotplots of Hi-C pseudo-molecules against HiFiasm contigs (Longest contigs >1Mb). (b) Dotplots of Hi-C pseudo-molecules against HiFiasm contigs (Longest and Middle size contigs). Red star showed the extra middle size contig (Table S2, 4^th^ column)

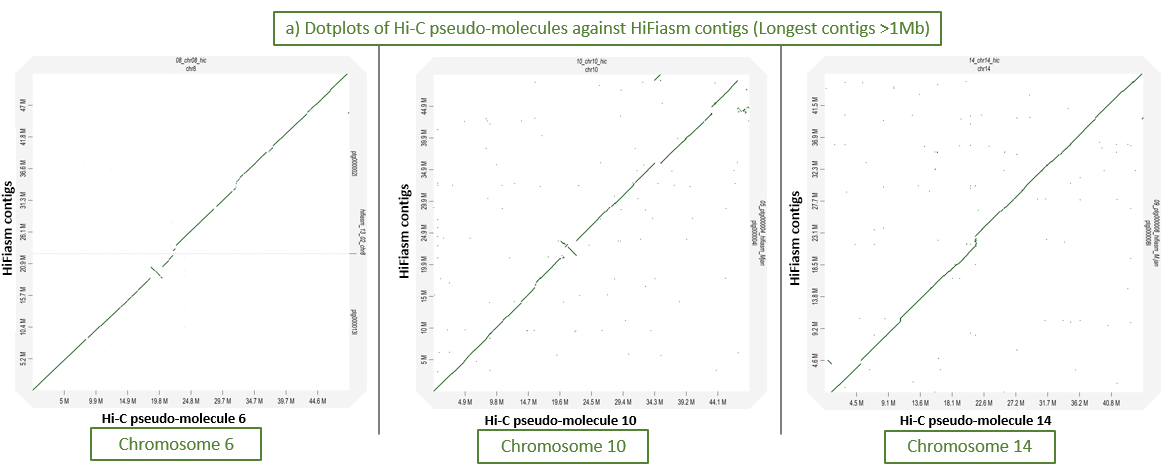

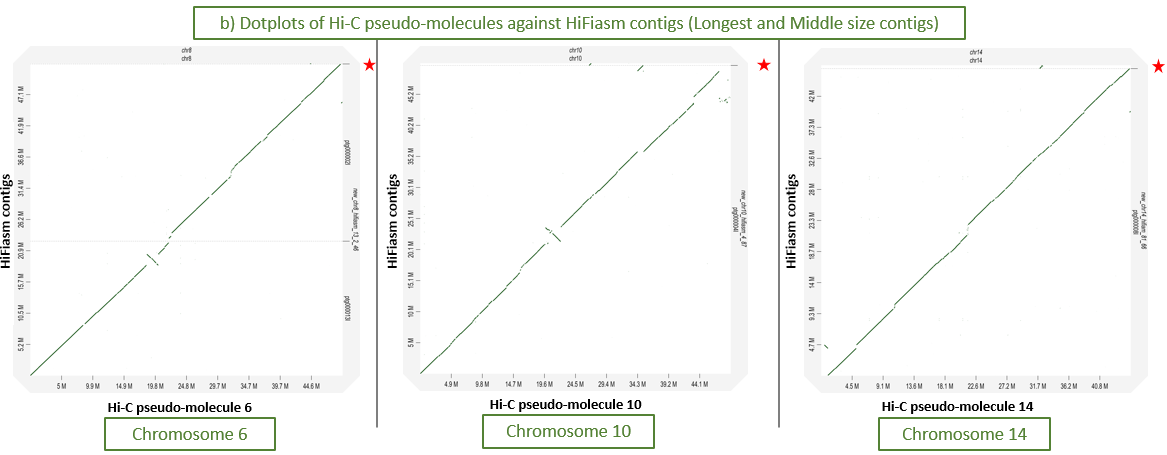

**Figure S3:** (a) Dotplots of Hi-C pseudo-molecules against HiFiasm contigs (Longest contigs >1Mb). S2 (b) Dotplots of Hi-C pseudo-molecules against HiFiasm contigs (Longest and Middle size contigs).

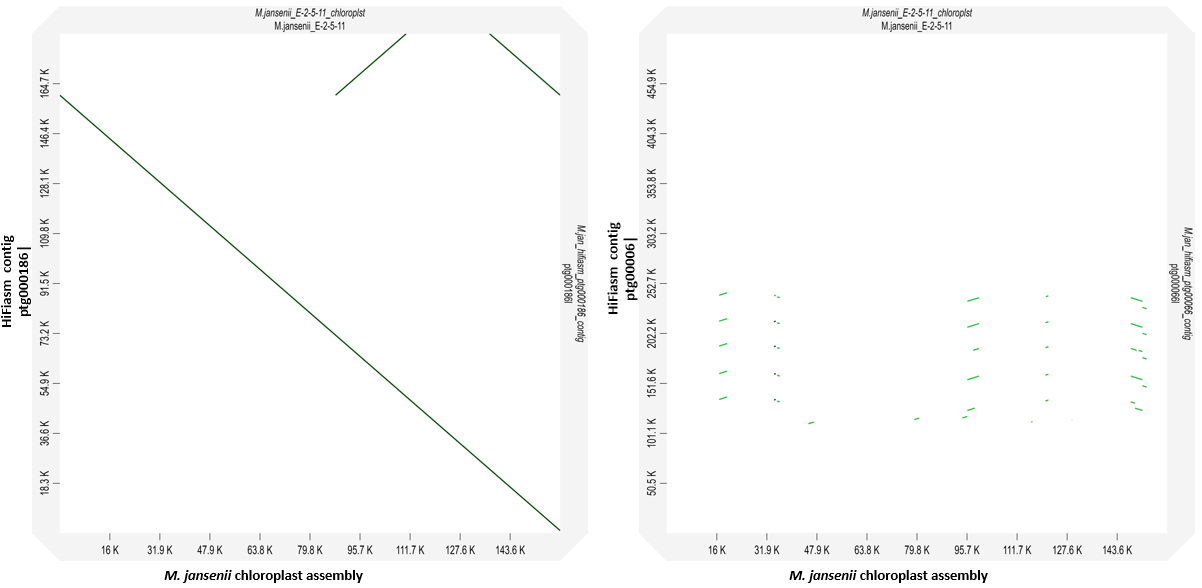

**Figure S4: Chloroplast assembly covered by single HiFiasm Contig (Ptg0000186) and small bits by Ptg000066.**

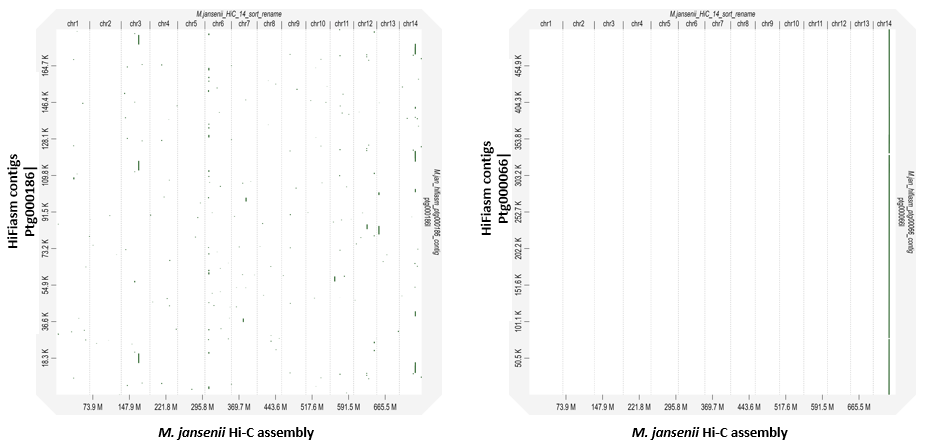

**Figure S5: Chloroplast sequence (Ptg0000186 & Ptg000066) insertions in the Hi-C assembly**

**Table S1: HiFiasm Contigs (separate excel file)**

**Table S2:** HiFiasm contigs (< 1Mb and > 100 Kb) which are part of Hi-C pseudo-molecule assembly.

***All middle size contigs (#64) of HiFiasm assembly corresponds to Hi-C assembly, but only contigs with “Alignment block length” greater than 100 Kb were considered.**

| **HiFiasm middle size contigs *** | **Length of contig** | **start** | **end** | **Strand** | **Hi-C chromosome** | **Length of chromosome** | **Start** | **end** | **Number of residue matches** | **Alignment block length** |
| --- | --- | --- | --- | --- | --- | --- | --- | --- | --- | --- |
| ptg000087l | 353246 | 6 | 333848 | + | chr10 | 48974653 | 26634412 | 26968299 | 319711 | 333951 |
| ptg000051l | 823842 | 510970 | 823838 | + | chr7 | 52077970 | 7635725 | 7942300 | 283252 | 314722 |
| ptg000109l | 388144 | 17 | 275074 | + | chr3 | 58143993 | 26511644 | 26786699 | 261393 | 275068 |
| ptg000099l | 388484 | 5047 | 272569 | + | chr5 | 55220784 | 26342363 | 26609914 | 259802 | 267725 |
| ptg000066l | 505394 | 92060 | 306715 | + | chr14 | 45288529 | 32032742 | 32247416 | 206578 | 214723 |
| ptg000071l | 117037 | 8 | 117031 | + | chr3 | 58143993 | 39146002 | 39262602 | 114711 | 117144 |
| ptg000231l | 173643 | 3 | 154150 | + | chr6 | 53595462 | 45346182 | 45497608 | 119329 | 166044 |
| ptg000046l | 103639 | 2 | 103632 | + | chr8 | 49563658 | 44424231 | 44527861 | 101007 | 103640 |

**Table S3: HiFiasm contigs (biggest contigs & middle size contigs) corresponds to M. jansenii Hi-C 14 pseudo-molecules.**

| ***M. jansenii* Hi-C pseudo-molecules** | **Size of Hi-C Pseudo-molecules** | **Hifiasm Contigs Corresponding to Hi-C scaffolds (> 1Mb)** | **Hifiasm Contigs Corresponding to Hi-C scaffolds (Middle_size contigs)** |
| --- | --- | --- | --- |
| Chr 1 | **67682215** | Ptg000016l (complete) | NA |
| Chr 2 | **63669590** | Ptg000006I (start small) + Ptg000025I + Ptg000010I (end large) | NA |
| Chr 3 | **58143993** | Ptg000021I (start small) + ptg000009l (middle large) + ptg000022I (end small) | ptg000109l + ptg000071l |
| Chr 4 | **56076407** | Ptg000017l (complete) | NA |
| Chr 5 | **55220784** | Ptg000011l (complete) | ptg000099l |
| Chr 6 | **53595462** | Ptg000003l (complete- fragmented) | ptg000231l |
| Chr 7 | **52077970** | Ptg000019I (start small) + Ptg000020I (middle) + ptg000033l + ptg000007I (last) | ptg000051l |
| Chr 8 | **49563658** | ptg000013I (start part) + ptg000002l (end part) | ptg000046l |
| Chr 9 | **49085581** | ptg000014l (start big) + ptg000005 (small end) | NA |
| Chr 10 | **48974653** | ptg000004l (complete but fragmented) | ptg000087l |
| Chr 11 | **47698009** | ptg000012l (complete) | NA |
| Chr 12 | **46713600** | ptg000001l (start half) + ptg000072I (middle tiny) + ptg000052I | NA |
| Chr 13 | **45610911** | ptg000023l (complete) | NA |
| Chr 14 | **45288529** | ptg000008l (complete) | ptg000066l |

**Table S4: IPA and HiFiasm assembly from different volumes of sequence data**

| **PacBio Sequence File details** | **Genome coverage** | **Assembler** | **Total number of contigs** | **Minimum length** | **Maximum length** | **Average length** | **N50** | **N75** | **Total length of assembly** | **BUSCO** |
| --- | --- | --- | --- | --- | --- | --- | --- | --- | --- | --- |
| CCS run 1 | 10 X | IPA | 1870 | 16 Kb | 3.5 Mb | 0.39 Mb | 0.61 Mb | 0.32 Mb | 736 Mb | 96.80% |
|  |  | HiFiasm | 4511 | 13 Kb | 7.2 Mb | 0.20 Mb | 0.38 Mb | 0.18 Mb | 909 Mb | 96.30% |
| CCS run 2 | 18 X | IPA | 728 | 13.7 Kb | 11.5 Mb | 1.00 Mb | 1.7 Mb | 0.99 Mb | 735 Mb | 95.70% |
|  |  | HiFiasm | 1058 | 15 Kb | 39.5 Mb | 0.79 Mb | 4.4 Mb | 2.1 Mb | 833 Mb | 98.40% |
| CCS run 1 + run 2 | 28 X | IPA | 284 | 11.4 Kb | 16.6 Mb | 2.6 Mb | 4.5 Mb | 2.6 Mb | 738 Mb | 98.00% |
|  |  | HiFiasm | 779 | 14.8 Kb | 71.9 Mb | 1 Mb | 46.1 Mb | 25.1 Mb | 826.7 Mb | 99.60% |
